# Cis-regulatory effect of HPV integration is constrained by host chromatin architecture in cervical cancers

**DOI:** 10.1101/2022.11.28.518229

**Authors:** Anurag Kumar Singh, Kaivalya Walavalkar, Daniele Tavernari, Giovanni Ciriello, Dimple Notani, Radhakrishnan Sabarinathan

## Abstract

Human papillomavirus (HPV) infections are the primary drivers of cervical cancers, and often the HPV DNA gets integrated into the host genome. Although the oncogenic impact of HPV encoded genes (such as E6/E7) is well known, the cis-regulatory effect of integrated HPV DNA on host chromatin structure and gene regulation remains less understood. Here, we investigate the genome-wide patterns of HPV integrations and associated host gene expression changes in the context of chromatin states and topologically associating domains (TADs). We find that HPV integrations are significantly enriched and depleted in active and inactive chromatin regions, respectively. Interestingly, regardless of the chromatin state, the genomic regions flanking HPV integrations showed transcriptional upregulation. Nevertheless, the upregulation (both local and long-range) is mostly confined to the TADs with integration and does not affect the adjacent TADs. Few TADs showed recurrent integrations associated with overexpression of oncogenes within them (such as *MYC, PVT1, TP63* and *ERBB2*), regardless of the proximity. To further understand the long-range effect, we performed HiC and 4C-seq analyses in HeLa and observed chromatin looping interaction between the integrated HPV and *MYC*/*PVT1* regions (situated ∼500 kb apart), leading to allele-specific overexpression of these genes. Again, these chromatin interactions involving integrated HPV are mostly observed within the same TAD. Together, these results suggest the cis-regulatory potential of integrated HPV DNA that drives host gene upregulation at intra-TAD level in cervical cancer. Based on this, we propose HPV integrations can trigger multimodal oncogenic activation to promote cancer progression.

## Introduction

Cervical cancer is the fourth most common cancer type among women worldwide, with the majority being reported from developing and underdeveloped countries (Sung et al. 2021). Studies have established that most cervical cancer cases can be attributed to the persistent Human papillomavirus (HPV) infection, particularly the high-risk subtypes such as HPV16 and HPV18 (Bhatla et al. 2008; Moody and Laimins 2010). HPV contains a circular double-stranded DNA genome of size ∼8 kb and it infects basal epithelial cells in the cervix. During the initial stages of infection, the HPV DNA exists in episomal form. However, over the course of epithelial cell differentiation, proliferation or neoplastic changes, the HPV DNA gets integrated into the host genome. This integration process is likely to occur at the host genomic regions sensitive to DNA strand breaks and share microhomology with the HPV DNA (Warburton et al. 2021b; Hu et al. 2015). Large-scale genome study of cervical tumours has shown that 80% of the tumours with HPV had integration in the host genome (Cancer Genome Atlas Research Network et al. 2017).

Tumours with HPV DNA integration show overexpression of viral oncogenes such as E6 and E7, likely due to the perturbations or DNA breakpoints in the viral regulatory gene E2 (which controls the expression of E6/E7) or increased stability of the viral transcripts upon fusion with the host genes (Moody and Laimins 2010). E6 and E7 proteins are known to interfere with the host p53 and RB pathways respectively, and thus favour cancer cell proliferation by avoiding apoptosis and cell cycle arrest (Yim and Park 2005; Hoppe-Seyler et al. 2018). Besides our understanding of the oncogenic roles of these viral proteins (E6/E7), efforts to delineate the cis-regulatory effects of HPV integration on the host chromatin structure and gene regulation have been rather limited.

HPV integration in the host genome can be a single or clustered (multiple nearby) event. The latter is often found together with genomic alterations (including amplification, deletion and translocations) in the nearby host regions, likely due to the HPV integration mediated DNA replication and recombination processes (Akagi et al. 2014; McBride and Warburton 2017). Besides, HPV integrations are associated with the upregulation of host genes which are either directly affected by the integration or in its immediate vicinity (Nguyen et al. 2018; Cancer Genome Atlas Research Network et al. 2017; Tang et al. 2013; Ojesina et al. 2014). Furthermore, recent studies using cell lines showed that HPV integration can cause long-range effects in *cis* through changes in the host chromatin interactions and subsequent gene dysregulation. For example, HPV16 integration in the W12 (human cervical keratinocyte) cell line was shown to alter chromatin interactions (involving both host:host and host:viral DNA), as well as host gene expression in the nearby regions (Groves et al. 2021). Similarly, in the cervical adenocarcinoma cell line HeLa, the HPV18 DNA integration in chromosome 8 was shown to have long-range chromatin interactions with the promoter region of the *MYC* oncogene (located approximately 500 kb away in the same chromosome) and was associated with its overexpression (Shen et al. 2017; Adey et al. 2013). However, the extent of these long-range chromatin interactions mediated by HPV integrations genome-wide and the associated host gene expression changes are still unexplored in cervical tumours.

Previous studies have shown that the HPV integrations from the cervical tumours and cell lines were enriched in the transcriptionally active open-chromatin regions (Doolittle-Hall et al. 2015; Bodelon et al. 2016; Christiansen et al. 2015). However, these findings were based on the HPV integrations derived mostly from transcription based assays (such as RNA-seq and amplification of papillomavirus oncogene transcripts (APOT)), and thus probably have a bias towards transcriptionally active regions. Hence, a whole-genome DNA based HPV integration detection approach is required to understand the distribution of HPV integrations across the genome and to study their impact on chromatin structure and gene-regulation. Moreover, a recent PCAWG consortium study has also demonstrated the need for DNA based methods to obtain a comprehensive view of viral association with cancers (Zapatka et al. 2020).

To address the aforementioned limitations, we explored the genome-wide HPV integration patterns and their impact on host gene expression in the context of chromatin states and topologically associating domains (TADs). For this, we collated genome-wide HPV integrations in cervical cancers detected using DNA based approaches and compared it with the chromatin state information from HeLa (cervical adenocarcinoma) and NHEK (normal human epidermal keratinocytes) cell lines. We found that the HPV integrations are significantly enriched in active chromatin regions and depleted in inactive chromatin regions, as compared to the expected counts. Interestingly, regardless of the host chromatin state, transcriptional upregulation was observed in the immediate vicinity of the HPV integration regions (up to 10 kb). Further investigation of the long-range effects of HPV integration revealed that the TADs with integration have higher gene expression as compared to samples without integration in the same TAD. More importantly, this difference was not observed in the TADs adjacent to the HPV integrated TADs. Moreover, the recurrent HPV integration analysis at the TAD level revealed both the direct and long-range effect of HPV integration on the expression of cancer related genes (such as *MYC, PVT1, TP63* and *ERBB2*). Additionally, we used Hi-C and 4C-seq analyses to show that the HPV integration in HeLa cells mediates long-range chromatin interactions with the oncogenes *MYC* and *PVT1*, and drives their overexpression in an allele-specific manner. Interestingly, these chromatin interactions were also mostly confined to the same TAD and not extending to the neighbouring TADs. Together, our results suggest the cis-regulatory potential of integrated HPV DNA that drives upregulation of host genes through changes in chromatin interactions (but mostly within the same TAD) in cervical cancer. This underscores that the HPV integration can mediate multiple modes of oncogenic activation and thereby acts as a strong driver conferring selective growth advantage to the cancer cells.

## Results

### HPV integrations are enriched in active chromatin regions and depleted in inactive chromatin regions

At first, we asked if the HPV integrations (n=617 from 212 samples) detected from the genome-wide DNA based approaches are distributed randomly or enriched in specific functional regions of the host genome. For this, we compared the HPV integration sites with ChromHMM annotations (Ernst and Kellis 2012; Roadmap Epigenomics Consortium et al. 2015), which categorise the genome into broad functional annotations based on various histone modification profiles, from two closest cell lines available: HeLa (cervical adenocarcinoma) and NHEK (normal human epidermal keratinocytes). To check for the enrichment, we compared the number of observed HPV integrations overlapping with each of these annotations with the expected counts computed using random sites of equal size and similar GC content (see Methods).

With both HeLa and NHEK ChromHMM annotations, we observed that compared to the expected, HPV integrations were significantly enriched (Chi-squared test, FDR<0.05) in the transcriptionally active regions (TxFlnk, Tx, TxWk), enhancers (EnhG, Enh), and zinc finger protein gene/repeat regions (ZNF/Rpts); whereas a significant depletion (FDR<0.05) was observed in polycomb repressed/heterochromatin regions (ReprPC, ReprPCWk, Het), and quiescent regions (in NHEK but not in HeLa) (Figure 1A, B). We further merged all the ChromHMM annotations into two major categories – active and inactive – based on the gene-regulation/chromatin activity of the regions (Roadmap Epigenomics Consortium et al. 2015) (see Methods). In both HeLa (Figure 1C) and NHEK (Figure 1D), HPV integrations were significantly enriched in active regions (one-sample Chi-squared test, P < 0.0001) and significantly depleted in inactive regions (P<0.05) as compared to the expected counts.

**Figure 1:**
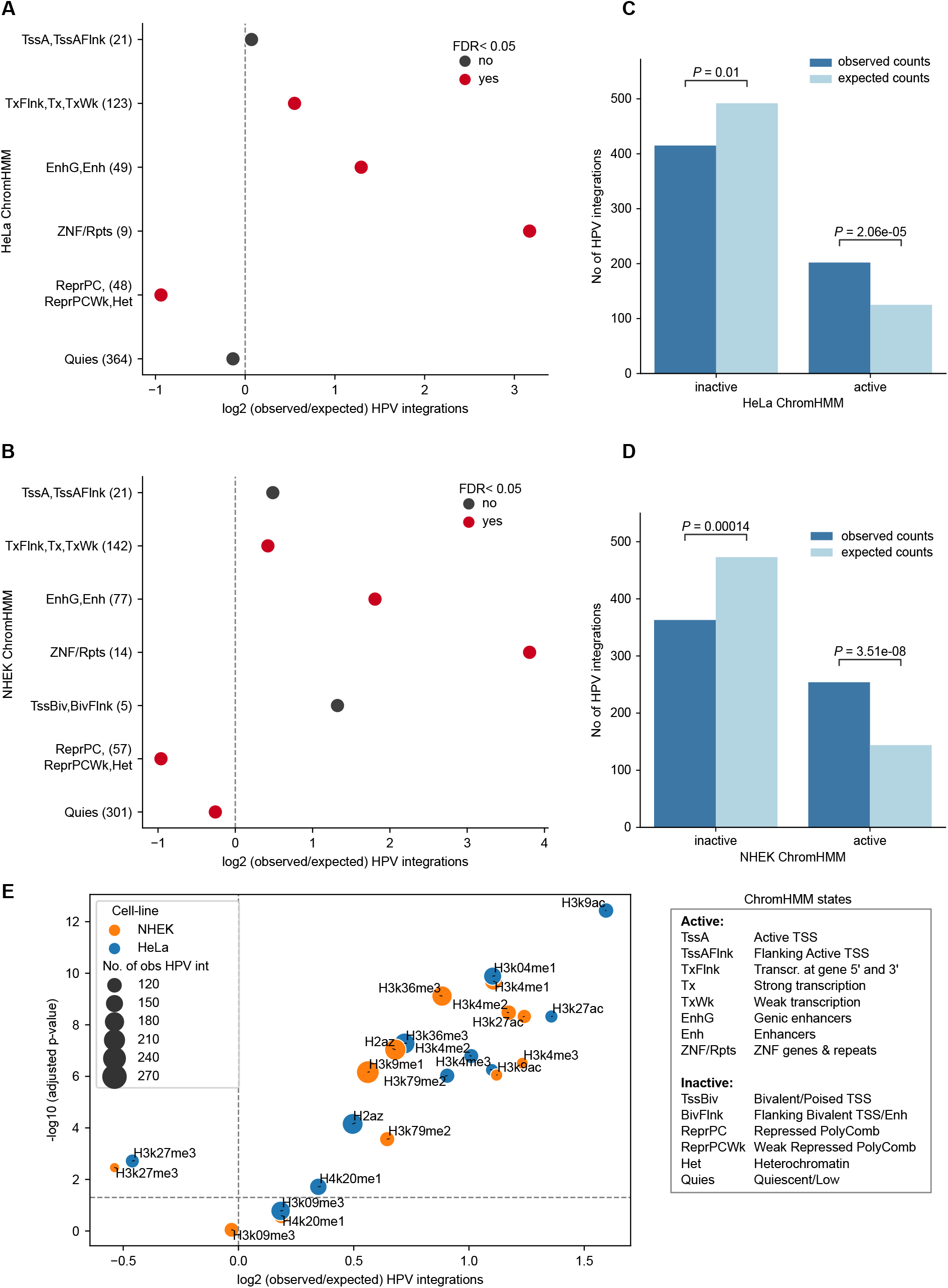
Enrichment of HPV integrations in functionally annotated regions. **A)** Enrichment of HPV integrations in the ChromHMM annotated regions from the HeLa cell line. The x-axis represents the log2 of observed/expected number of HPV integrations. The y-axis represents the different annotations and the observed number of HPV integrations overlapping them (given in the bracket). The p-values were computed using Chi-squared goodness-of-fit test followed by FDR correction. The colour of the dots indicates whether the adjusted p-value is below the significance level of 5% or not. **B)** Same as **(A)** but with ChromHMM annotations from the NHEK cell line. **C-D)** Bar plot showing the frequency of observed and expected integrations in the active and inactive regions defined by combining ChromHMM annotations in HeLa **(C)** and NHEK **(D)** (see Methods). The p-value was calculated using a one-sample Chi-squared test. **E)** Enrichment of HPV integration in various histone modification regions from HeLa and NHEK. The x-axis represents the log2 of observed/expected number of HPV integrations and the y-axis represents the negative log10 of adjusted p-value (Chi-squared test followed by FDR correction). The horizontal dashed line represents FDR cut-off of 5%. The colour and size of the dots represent the cell line and the number of observed HPV integrations for each of the histone marks, respectively.

Further, we checked specifically which histone modification marks associated with the above annotations are enriched with HPV integrations. We observed that the HPV integrations were significantly enriched (Chi-squared test, FDR<0.05) in various active histone modification regions (such as H3K4me1, H3K4me2, H3K4me3, and H3K27ac) in both HeLa and NHEK (Figure 1E). In contrast, HPV integrations were significantly depleted (FDR<0.05) only in the repressive histone modification regions (H3K27me3) in both HeLa and NHEK (Figure 1E). Despite these two cell lines being different (HeLa-malignant versus NHEK-non malignant), the HPV integration patterns remain consistent. This suggests that the host cell-type specific chromatin structure influences the HPV integration patterns regardless of malignancy.

HPV integration has previously been shown to be enriched in regions of fragile sites (Warburton et al. 2021b), which contain DNA repeats that could form non-B DNA conformations and are associated with genomic instability. To check this further, we computed the enrichment of HPV integrations in the DNA regions predicted to form non-B DNA conformations (see Methods). Among all the non-B forms of DNA, only direct repeats showed significant enrichment of HPV integration as compared to the expected (Chi-squared test, FDR<0.05) (Supp Figure 1A). Further, to check this in the context of chromatin states, we calculated the odds ratio of observed versus expected HPV integrations in active versus inactive regions (for HeLa and NHEK, separately). This showed that even in the non-B DNA regions, integration tends to occur more frequently in the active regions as compared to inactive regions (odds ratio > 1 and Fisher exact test, FDR < 0.05), except for A phased repeats (with both HeLa and NHEK ChromHMM) and G quadruplex regions (only with HeLa ChromHMM) (Supp Figure 1B, C).

Taken together, these results suggest that HPV integrations are not randomly distributed across the genome. They are highly enriched in the active chromatin regions and depleted in repressed chromatin regions. This could be due to the combined or individual effect of DNA sequence context, DNA accessibility, DNA damage response (linked to the chromatin states) and positive selection.

### HPV integration affects host transcriptional activity regardless of the host chromatin state

Previous studies have shown that HPV integration can affect host gene expression in its immediate vicinity (Nguyen et al. 2018; Ojesina et al. 2014). We asked if this effect could be influenced by the host chromatin states. To test this, we used TCGA cervical cancer (CESC) samples (only WGS with matched gene expression data available, n=50) with 151 HPV integration regions and plotted the total RNA expression (both genic and non-genic elements) in the 10 kb flanking region around the integration (Nguyen et al. 2018) (see Methods). We observed that the samples with HPV integration showed increased expression in their neighbouring 10 kb regions as compared to mean expression from the samples without HPV integration in the same region, regardless of whether the HPV integration was in an active (Mann-Whitney U test, P=0.0048 for HeLa and P=7.28e-05 for NHEK) or inactive (P=0.0065 for HeLa and P=0.057 for NHEK) region (Figure 2A, Supp Figure 2A). Following that, we asked whether the activity of regulatory elements (like enhancers) are also enhanced around the HPV integration regions. To test this, we plotted enhancer RNA or eRNA expression (indicative of the functional activity of enhancers) from the super enhancer (SE) regions, within 10 kb on either side of the HPV integration regions in the above TCGA-CESC samples (Figure 2B, Supp Figure 2B). We observed an increased eRNA expression in the HPV integrated samples compared to the samples without integrations. Again as shown above, it was not influenced by the chromatin state of the integrated region.

**Figure 2:**
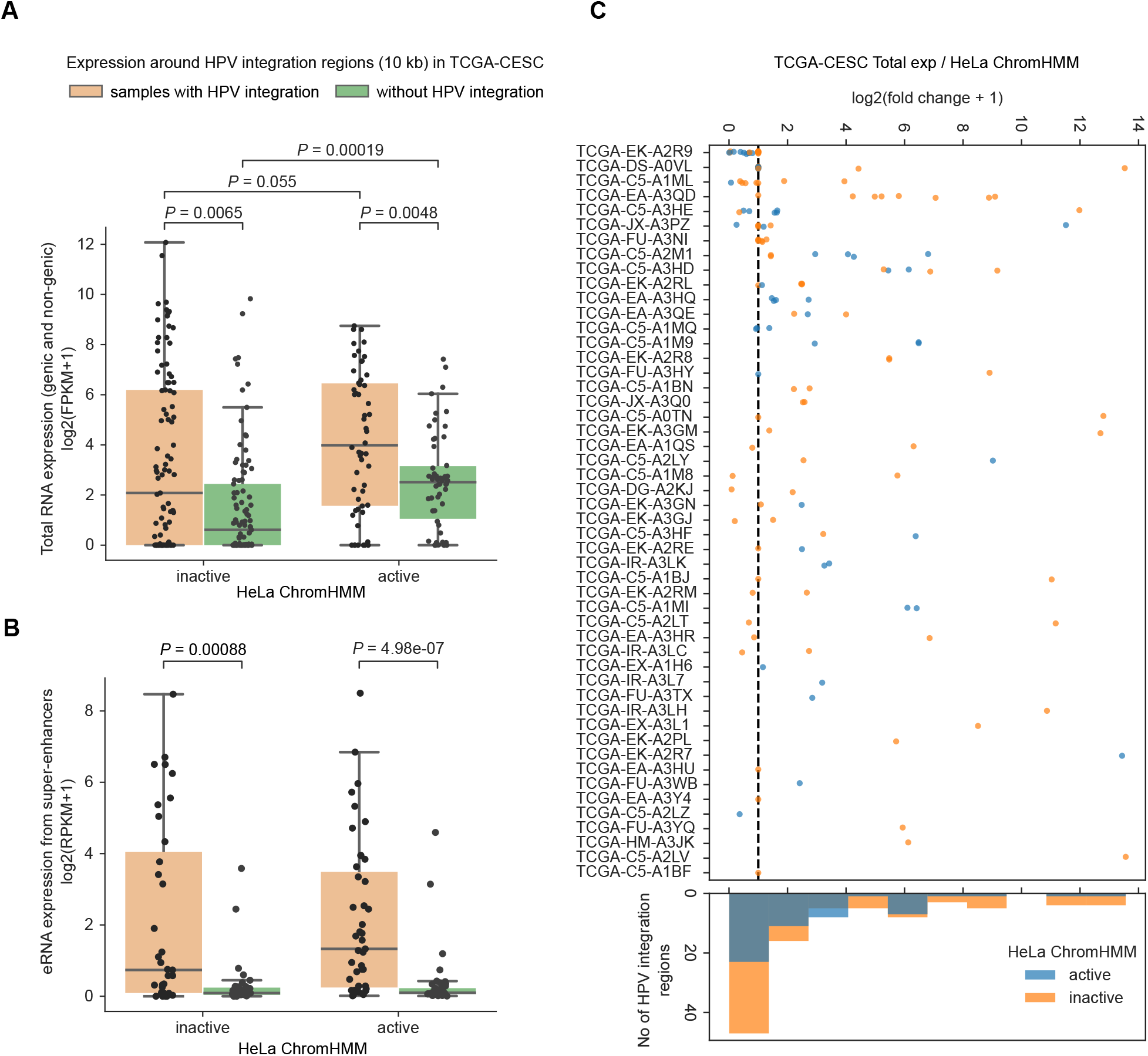
Enhanced transcriptional activity near HPV integration in the context of chromatin states from HeLa. **A)** Boxplot showing the total expression in the 10 kb flanking region around the HPV integration regions as compared to mean expression from TCGA-CESC samples without HPV integration in the same region. The x-axis represents whether the HPV integration is located in an active or inactive chromatin region with respect to HeLa ChromHMM. The p-values were computed using Mann-Whitney U test (two-sided). **B)** Same as **(A)** but for the eRNA expression from super enhancers within 10 kb on either side of the HPV integration regions. **C)** Expression fold change associated with each of the HPV integration regions. The x-axis represents the log2 fold change, which was calculated as the total expression in the 10 kb flanking region around HPV integration regions divided by the mean expression from other samples without HPV integration in the same genomic region. The y-axis represents the individual sample-id of TCGA-CESC samples. The colour of the dots indicates if the integration overlaps an active or inactive ChromHMM region of HeLa. The black vertical line represents the value of log2(fc+1)=1. The histogram at the bottom shows the frequency of integration regions at different fold-change bins. Each dot in **(A-C)** represents a HPV integration region from a sample.

We additionally noticed that the control samples (without HPV integration) showed significantly higher expression in active as compared to inactive regions (P=0.00019 for HeLa, Figure 2A and P=0.00031 for NHEK, Supp Figure 2A). This underscores that the ChromHMM annotations from cell lines (HeLa/NHEK) match well with the tumour tissues in terms of transcriptional activity observed in the active and inactive regions. More importantly, in samples with HPV integration, we observed a higher expression in the active as compared to the inactive regions (P=0.055 for HeLa, Figure 2A and P=0.0061 for NHEK, Supp Figure 2A). This indicates that the HPV integration in active chromatin regions further enhances the host transcription activity in its vicinity. Further, we asked whether all HPV integrations in a sample could lead to overexpression in its immediate vicinity or not. For this, we computed the expression fold change for each HPV integration region (ratio of expression in the 10 kb flanks around integration regions with the mean expression from other samples without integration in the same region) (Nguyen et al. 2018). This showed that the expression association with HPV integration was highly variable and not all HPV integration regions associated with higher transcriptional activity in its vicinity (Figure 2C, Supp Figure 2C). This could be due to the epigenetic suppression of certain integrated HPV regions or impaired regulatory activity (Warburton et al. 2021a).

Taken together, these results suggest that the HPV integration leads to transcriptional upregulation and enhanced enhancer activity in its immediate vicinity, regardless of the host chromatin state. Nevertheless, the transcriptional activity associated with HPV integration was relatively higher in active chromatin regions, as compared to inactive regions.

### Transcriptional activity associated with HPV integration is mostly confined to the same TAD

We next asked whether the HPV integration can mediate or alter the host long-range chromatin interactions (such as enhancer-promoter or promoter-promoter), and thereby dysregulate gene expression in tumours. To test this, we compared the host gene expression at the level of TADs, obtained from HeLa and NHEK cell lines (Rao et al. 2014). TADs act as functional units of genome organisation by restricting the interactions between regulatory elements and thereby controlling gene regulation (Rao et al. 2014; Li et al. 2018). TAD boundaries are commonly bound by insulator proteins like CTCF that prevent interactions across the TADs. We hypothesised that the transcriptional overexpression associated with the HPV integration would be majorly restricted to the TADs where the HPV integration is localised. To test this, we plotted the average gene expression at the TAD level in the TCGA-CESC samples with HPV integration and compared it with the mean expression from samples without integration in the same TAD (see Methods). We further extended this analysis to the immediate upstream (5’) and downstream (3’) TADs for comparison. Only the TADs with HPV integration showed overall increased expression compared to the samples without integration (Wilcoxon signed-rank test, P<0.0001) (Figure 3A, Supp Figure 3A). However, neither the upstream nor the downstream neighbouring TADs showed any effect at TAD level expression of genes (P>0.01) (Figure 3A, Supp Figure 3A).

**Figure 3:**
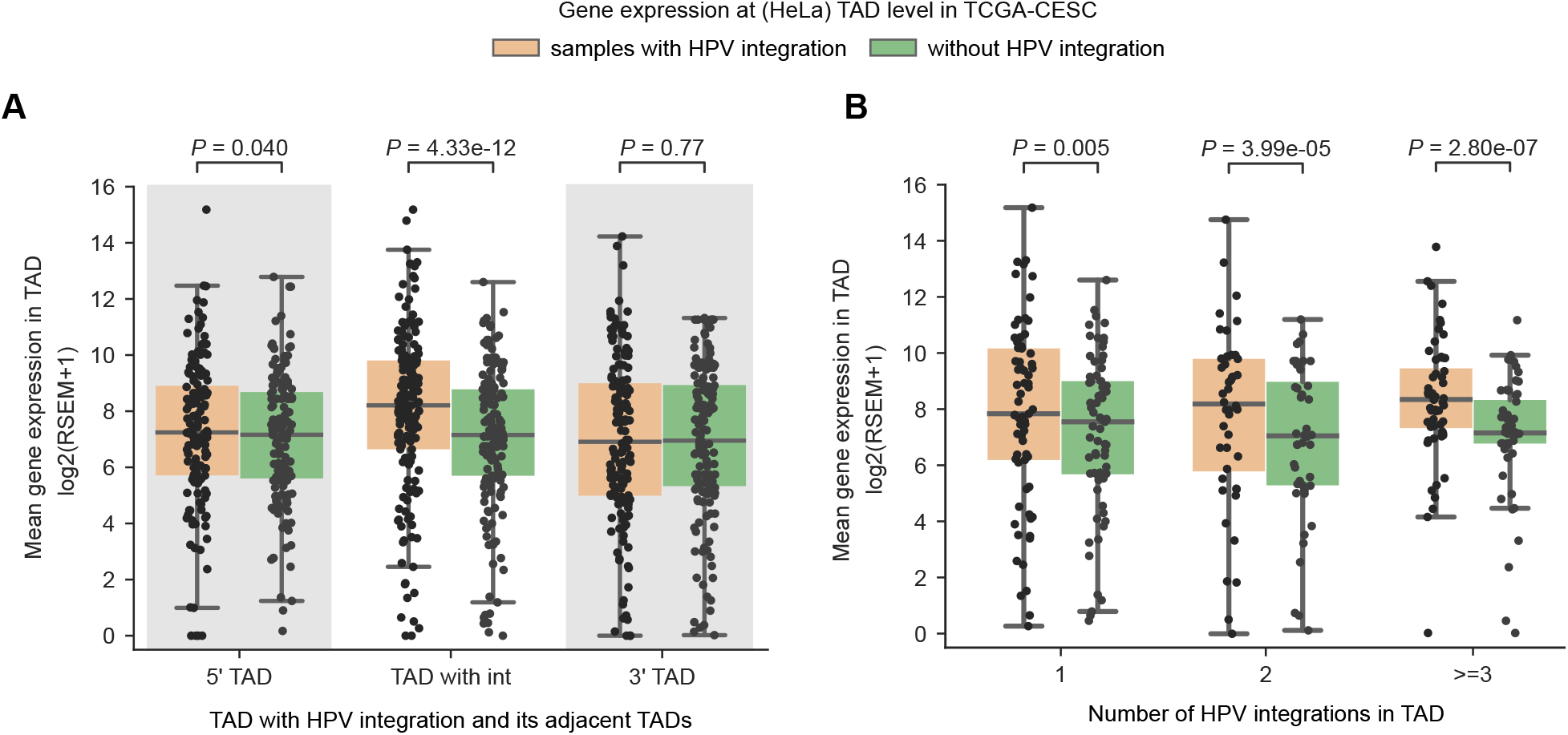
HPV integration associated host gene overexpression with respect to HeLa TAD domains. **A)** TAD level gene expression in the TCGA-CESC samples with HPV integration compared to the mean expression from the samples without HPV integration in the same TADs, also for the neighbouring upstream (5’) and downstream (3’) TADs. The TAD information was obtained from the HeLa cell line. **B)** TAD level gene expression in the TCGA-CESC samples with HPV integration, separated by whether the TAD had 1, 2 or more than 2 integrations compared to the mean expression from the samples without HPV integration in the same TADs. Each dot in **(A)** and **(B)** represents a unique HeLa TAD-tumour sample combination and the p-values were computed using the Wilcoxon signed-rank test (two-sided).

Further, we asked whether increase in the number of integrations in a TAD would result in more perturbations in gene regulation and overexpression. For this, we separated TADs based on whether they had 1, 2, or more than 2 integrations and compared the TAD level gene expression among them, along with the samples without integration. This showed that indeed an increase in the number of integrations in a TAD was associated with an increase in gene overexpression as compared to the samples without integration in the same TADs (Figure 3B).

Direct HPV integration in the host genes (or fusions) can lead to overexpression of the target genes. To remove the influence of these, we repeated the above analysis after removing the genes which were directly overlapping with the HPV integration sites (Supp Figure 3B). Nevertheless, we found that the gene expression was higher in the TADs with integration as compared to samples without integration (P=3.36e-08), albeit slightly lower than the above (Figure 3A). This trend was consistent even after removing the genes which were either directly overlapping an integration or within 10 kb region flanking the integration (P=2.22e-07) (Supp Figure 3C), suggesting the long-range cis-regulatory potential of the HPV integration.

Taken together, these results indicate the potential of HPV integration to enhance expression of nearby host genes but mostly within the same TAD, likely due to the constraint imposed by the TAD boundaries or genome organisation. Further, within the TAD, the overexpression observed could come from both the direct and long-range effect of HPV integration on target genes through chromatin contacts in 3D nuclear space.

### Recurrent HPV integrations in TADs are associated with oncogene overexpression

Recurrent HPV integrations near cancer related genes (such as MYC and TP63) have been previously reported in cervical cancers (Warburton et al. 2021b; Hu et al. 2015). In those studies, the recurrence was defined mostly based on the HPV integration directly overlapping with or in close proximity (at a defined distance cut-off) of the target genes. This can potentially miss out the integrations that are far away and can have a long range effect on the target genes. To overcome this, we performed recurrent integration analysis at the TAD level. For this analysis, we considered HPV integrations (n=1324 from 326 samples) collated from both DNA-based studies (WGS, WXS and Hybrid capture) and RNA-seq (see Methods). The TADs which had HPV integration in at least three samples were considered recurrent (Figure 4A). This showed that TAD3189 and TAD2252 were the top two most recurrently integrated TADs. Interestingly, these TADs also exhibited mutually exclusive patterns of HPV integration, indicating that these recurrent TADs were not affected in the same patients. Moreover, we found that certain HPV subtypes were frequently integrated in specific TADs (Figure 4A). HPV16 was predominantly observed in TAD3189 (13/22), TAD2252 (11/15), TAD1333 (5/5) and TAD2320 (2/3), whereas HPV18 was predominant in TAD1369 (3/6). In the TAD3189, we observed both HPV16 and HPV18, however, the proportion of HPV18 was much higher (41%) as compared to the overall HPV18 proportion (16%) in TCGA-CESC (Chi-squared test, P=0.011).

**Figure 4:**
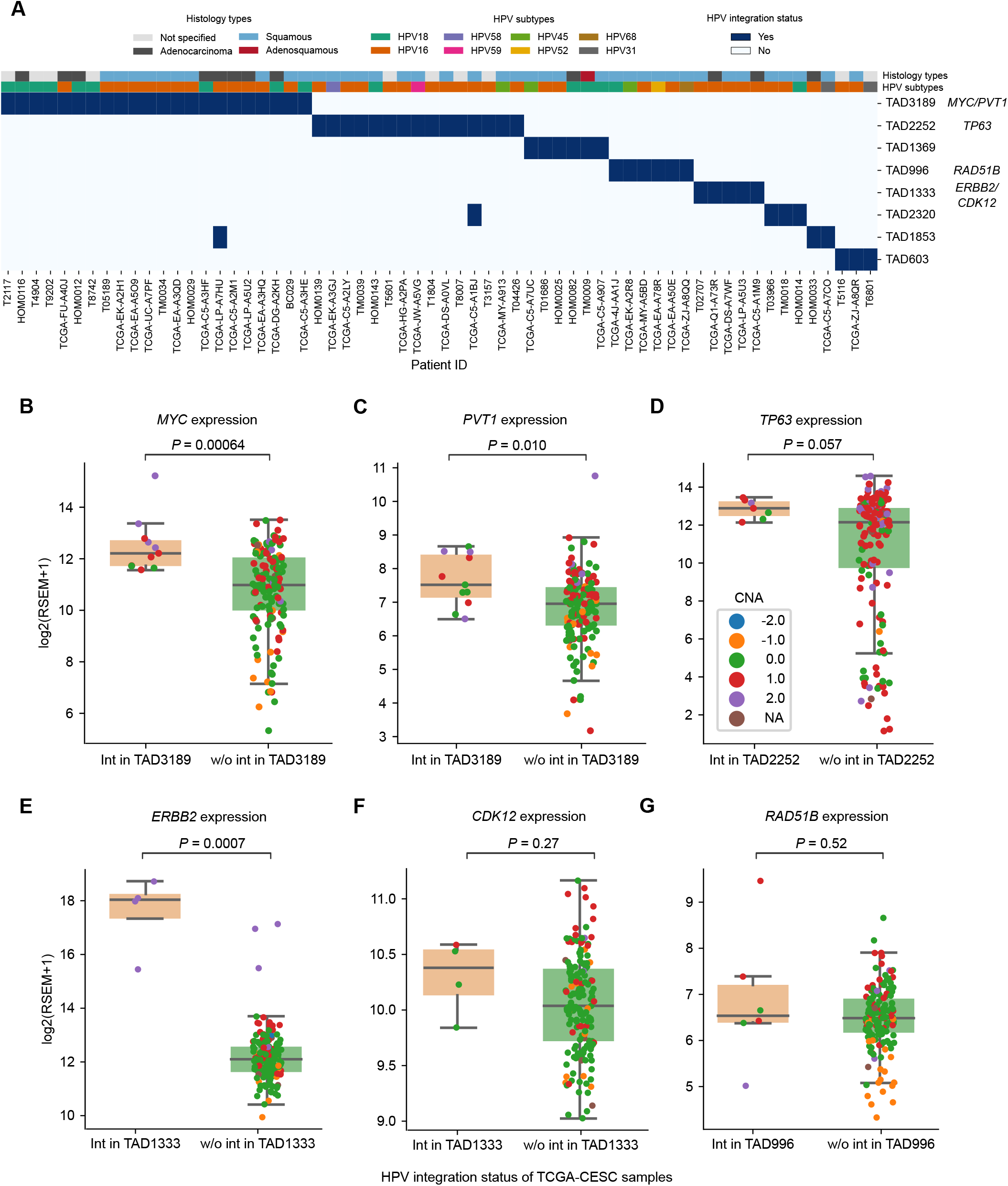
Recurrently integrated TADs and expression alteration of associated cancer genes. **A)** Heatmap shows the HeLa TADs with recurrent HPV integrations. The x-axis represents the sample-id and y-axis represents the TADs (denoted with distinct numbers to differentiate each TAD domain). Blue box indicates HPV integration in a particular TAD, and in a particular sample. The top two rows represent the tumour histology and HPV subtype in each of the samples, respectively. **B-C)** Gene expression of *MYC* **(B)** and *PVT1* **(C)** in TCGA-CESC samples with and without HPV integration in TAD3189. **D)** Gene expression of *TP63* in TCGA-CESC samples with and without HPV integration in TAD2252. **E-F**) Gene expression of *ERBB2* **(E)** and *CDK12* **(F)** in TCGA-CESC samples with and without HPV integration in TAD1333. **G**) Gene expression of *RAD51B* in TCGA-CESC samples with and without HPV integration in TAD996. The p-value shown in panels (**B-G**) was calculated using the Wilcoxon rank-sum test. The colour of each dot in the box plot (**B-G**) represents the relative copy number status of the gene in the respective TCGA-CESC samples (-2 deep deletion, -1 deletion, 0 copy neutral, 1 amplification, 2 high amplification).

To further understand if the recurrent HPV integrations are associated with tumorigenesis, we looked at the presence of cancer related genes (from Cancer Gene Census (Sondka et al. 2018) and Cancer LncRNA Census (Carlevaro-Fita et al. 2020)) in these TADs. Interestingly, TAD3189 harbours multiple important coding (*MYC*) and non-coding (*PVT1* and *CCAT1*) oncogenes. Overexpression of these oncogenes have been previously reported in cervical cancers and were also associated with poor prognosis (Narisawa-Saito et al. 2012; Iden et al. 2016; Shen et al. 2019). Thus, to test if HPV integration in TAD3189 is affecting the expression of these oncogenes, we divided the TCGA-CESC samples into two groups, one which had integration in TAD3189 and other which did not. Expression of both *MYC* (Wilcoxon rank-sum test, P=0.0006) (Figure 4B) and *PVT1* (P=0.010) (Figure 4C) was significantly higher in the samples with integration in TAD3189 as compared to samples without integration in TAD3189. Similarly in TAD2252, *TP63* (oncogene) expression was higher (P=0.057) (Figure 4D) in samples with integration in TAD2252. In TAD1333, *ERBB2* (oncogene) expression was significantly higher (P=0.0007) in samples with integration, whereas *CDK12* (tumour suppressor gene) in the same TAD did not show any change in expression (P=0.27) (Figure 4E, F). Also, *RAD51B* (tumour suppressor gene) in TAD996 did not show any change in expression level (P=0.52) (Figure 4G). This suggests that the HPV integration preferentially enhances the expression of oncogenes in these recurrent TADs. Interestingly, in these TADs (TAD3189 and TAD2252) with oncogenes, increased gene expression (z-score>0; above the average expression level across samples) was evident regardless of whether the HPV integration occurred directly at the gene or further away in the respective TADs (Supp Figure 4), suggesting that the HPV integration can affect gene expression both locally and at longer distances.

### HPV18 induced chromatin interactions in HeLa are confined locally and target oncogenes

Next, we asked whether the long distance effect of HPV integration on the oncogene expression in the above recurrent TADs could be mediated by chromatin looping. To study this, we chose the HeLa cell line (as a model) which has HPV18 integration in TAD3189 (chromosome 8), the most frequently HPV integrated TAD in cervical cancers (Figure 4A). To obtain an unbiased and global overview of all the genomic interactions between the integrated HPV DNA and the host genome, we leveraged the available Hi-C data from HeLa (Yardımcı et al. 2019), which captures overall genomic interaction at a particular resolution (or regions that are in close proximity in 3D space). For the Hi-C data analysis, we used the full length HPV18 genome (treated as a separate chromosome) along with human chromosomes, and mapped reads to compute the contact frequency (see Methods).

First, we looked at the chromosome level to identify which of the chromosomes have contacts with the integrated HPV18 DNA. This showed that the majority of the contacts involving HPV18 were associated with chromosome 8 (chromosome with HPV18 integration), followed by intra-HPV18 contacts (Figure 5A). Second, to gain further insights into these contacts with chromosome 8, we plotted the contact frequency (per kb) with the HPV18 for all the TADs on chromosome 8. This showed that the highest number of contacts with the integrated HPV18 DNA were observed with genomic regions in TAD3189, where the HPV integration localised (Figure 5B, Supp Figure 5A). Similarly, TAD3189 also had the highest coverage of reads (per kb) for the interaction between genomic regions on chromosome 8 and HPV18. This signifies that the interaction frequency between integrated HPV18 DNA and TAD3189 was much higher as compared to other TADs on the same chromosome (Supp Figure 5B, C). Together, these results demonstrate that the chromatin interactions driven by the integrated HPV18 are mostly confined to the same TAD containing the HPV18 integration, thus possibly supporting the TAD level upregulation of genes observed in tumours (see Figure 3A, Supp Figure 3A).

**Figure 5:**
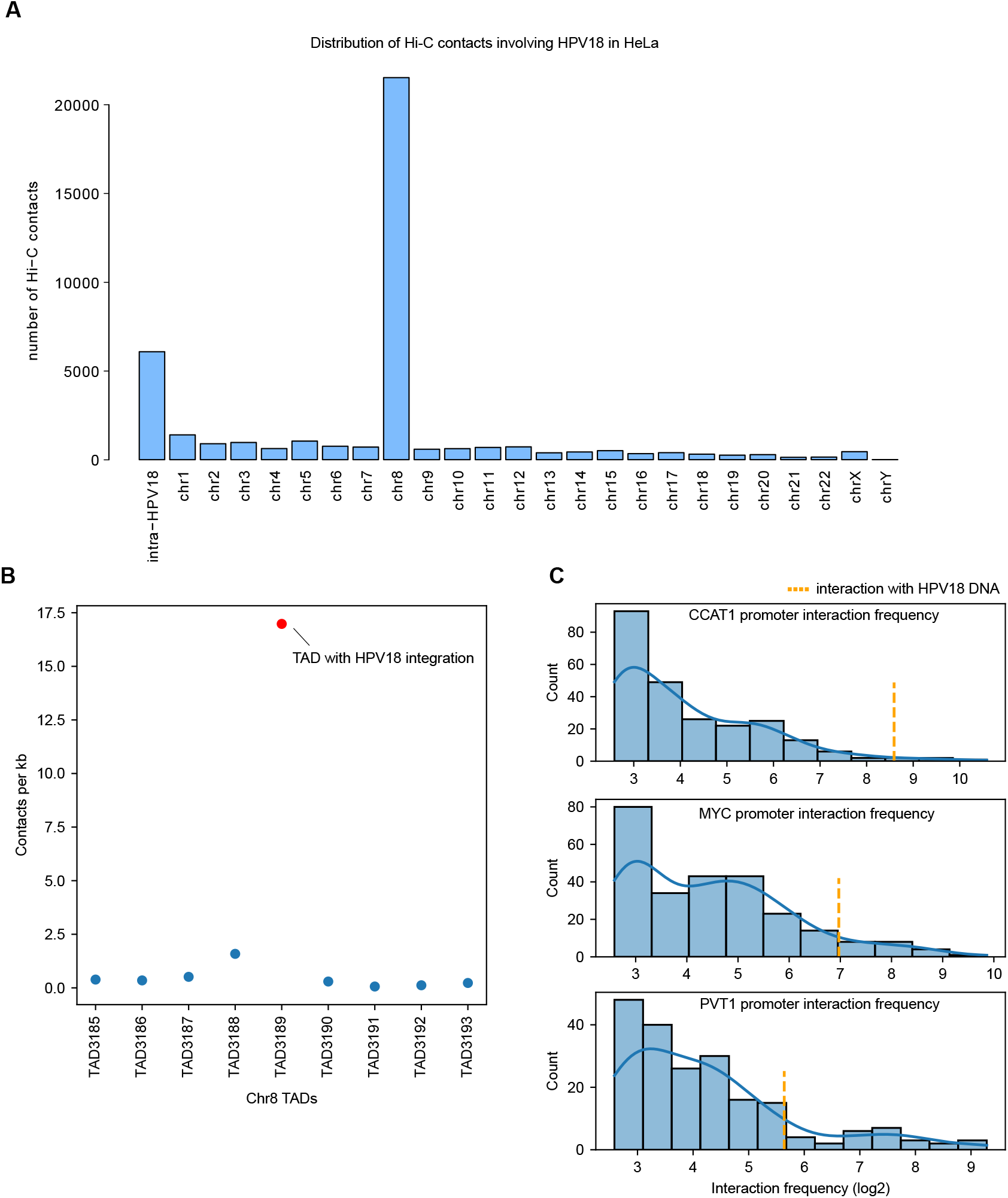
Hi-C analysis reveals highly localised chromatin interactions between integrated HPV18 DNA and host genome in HeLa. **A)** Bar plot shows the number of contacts between each of the chromosomes and the integrated HPV18 genome in HeLa, and also intra-HPV18 contacts. **B)** Scatterplot shows the contacts per kb between TADs on chromosome 8 and the HPV18 genome. All the TADs are arranged in a linear manner (5’ to 3’ direction of the genome). TAD3189 contains the HPV integration (red dot) which showed higher contacts as compared to the adjacent TADs (and also at the chromosome level, Supp Figure 5A). **C)** Histogram distribution plot shows the interaction frequency of the bins which overlap the promoters of *CCAT1, MYC* and *PVT1* with all the interacting bins on chromosome 8 and the integrated HPV18 (orange line). Only those interacting bins which were supported by more than 5 reads were shown in the plot.

We wanted to further understand the specific host regions which looped with the integrated HPV18 with a high frequency. For this, we selected the top 1% genomic bins (10 kb) from chromosome 8 with the highest frequency of interaction with HPV18, and overlapped them with the gene coordinates on the same chromosome. This resulted in the detection of 8 genes (*RP11-255B23.4, CCAT1, CASC8, RP11-382A18.3, RP11-382A18.2, CASC11, MYC, PVT1*) and all of them were located in the TAD3189 region. Of these, three are known oncogenes (*CCAT1, MYC* and *PVT1*). Further, we focused on the promoter region of these three oncogenes and observed that their interaction with HPV18 (among other regions on chromosome 8) was in top 1 percentile for *CCAT1* and *MYC* and top 2 percentile for *PVT1* (Figure 5C). This indicates a high frequency of chromatin interactions between these oncogenes and also their promoters with the integrated HPV18 DNA in HeLa.

### 4C-seq reveals haplotype-specific chromatin looping between integrated HPV18 and host genome

To further characterise the chromatin looping interaction mediated by the HPV18 integration at the local scale with higher resolution, we performed a 4C-seq experiment with the integrated HPV18 DNA as an anchor point in HeLa. At first, we wanted to check the extent of HPV integration induced chromatin interactions at the TAD level. For this, we plotted per bp coverage of 4C-seq reads in TAD3189 (with HPV integration) and in the neighbouring TADs (4 upstream and 4 downstream). Most of the 4C-seq signal was observed from the TAD with HPV integration similar to Hi-C analysis (Supp Figure 6A, B). Even though the HPV integration was found near the left boundary of TAD3189, the coverage of reads in the immediate upstream TAD was quite low. This suggests the role of chromatin structure (TAD boundaries) in confining the regulatory effects of the integrated HPV DNA within the same TAD predominantly. Further within the TAD3189, higher level of interaction was observed between the integrated HPV18 and all the three oncogenes (*CCAT1, MYC* and *PVT1*) in that TAD (Figure 6A, 4C track). This further supports the previous studies which showed an interaction between HPV18 and *MYC* locus using ChIA-PET (Adey et al. 2013) and 3C analysis (Shen et al. 2017).

**Figure 6:**
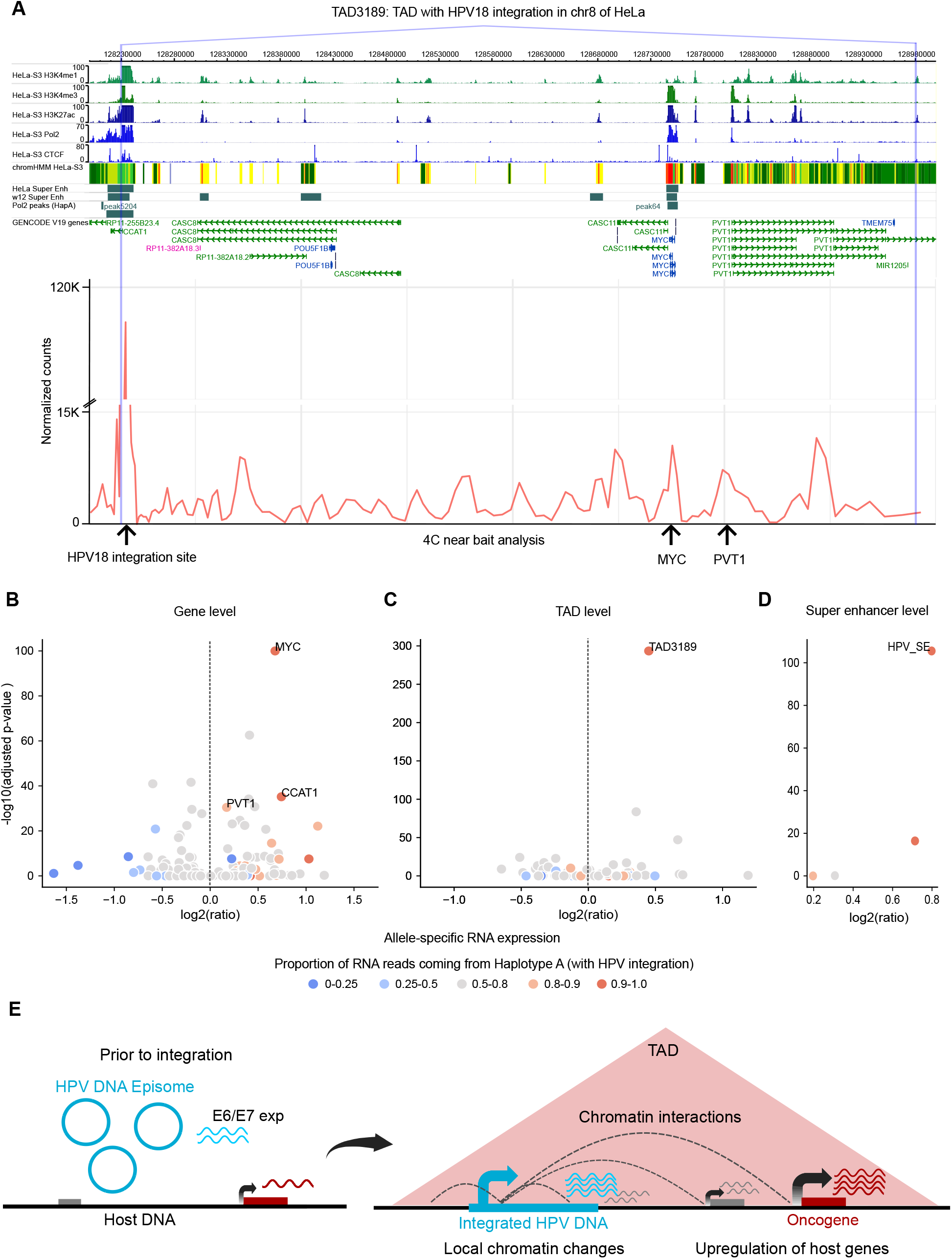
HPV18 integration mediated chromatin interactions from 4C-seq and allele-specific expression of oncogenes and SE in HeLa. **A)** Regions in the TAD3189 which show high interactions with the integrated HPV18 DNA by 4C-seq are shown in the bottom line plot (near bait analysis). The HPV integration region, *MYC* and *PVT1* is marked by an arrow. Various histone modifications, Pol2 and CTCF binding tracks from HeLa are also overlaid. Pol2 peaks (HapA) track shows the Pol2 peaks which were significantly haplotype specifically binding to Haplotype A. **B-D)** Allele-specific expression analysis for **B)** individual genes, **C)** TADs, and **D)** super-enhancers on chromosome 8. Values on the x-axis represent the log2 ratio of fraction of RNA and DNA reads coming from haplotype A (sum over all the heterozygous SNPs in a particular feature). The y-axis represents the negative log10 of Bonferroni corrected p values, after a binomial test of RNA read counts from Haplotype A against the proportion of DNA read counts from Haplotype A, for each of the features. **E)** Model summarising HPV integration mediated changes in host chromatin structure and gene expression dysregulation. HPV integration can lead to local chromatin changes resulting in transcriptional upregulation of host genes (also in fusion with viral genes) in its vicinity. In addition, the integrated HPV DNA can mediate long-range chromatin interactions resulting in the upregulation of genes at a distance.

In HeLa, integration of HPV18 is observed in only one of the two haplotypes of chromosome 8. Based on this, we hypothesised that the chromatin interaction observed between the integrated HPV18 DNA and the host genome could be haplotype-specific. To check this, we performed haplotype-specific 4C-seq coverage analysis, using heterozygous SNPs (as marker positions) from the HeLa genome (see Methods). This revealed that almost all reads from the 4C-seq (∼99-100%) mapped to the allele from the Haplotype A (which has the HPV integration) (Supp Figure 6C, D). This suggests that the HPV integration mediated chromatin interactions are not only localised mostly within the same TAD but are also haplotype specific.

### Haplotype-specific cis-regulating activity of HPV integration is associated with allele-specific oncogenes overexpression

Further, we asked if the above haplotype-specific chromatin looping interaction could lead to haplotype-specific regulatory changes and gene expression alterations. To check this, we performed haplotype-specific analysis for RNA polymerase II (Pol2) binding on chromosome 8 in HeLa. This resulted in three peaks of Pol2 in the TAD3189 which showed significant haplotype-specific binding to Haplotype A (>98% reads from Haplotype A in all 3), as compared to the expected proportion based on DNA copy number. Interestingly, these peaks overlapped with the *CCAT1* and *MYC* genes and also the super-enhancer overlapping the HPV integration (Figure 6A, Pol2 peaks (HapA) track), suggesting that the integrated HPV DNA drives gene regulation in haplotype specific manner. Further, we asked if this results in preferential expression of genes from the haplotype with HPV integration. To test this, we performed allele-specific gene expression analysis (taking into account the DNA copy number) on chromosome 8 in HeLa. This revealed *MYC* having significantly higher expression from Haplotype A (Figure 6B) (see Methods). The other oncogenic lncRNAs in the TAD3189, *CCAT1* and *PVT1*, also showed a similar pattern (Figure 6B). Further, we performed allele-specific gene expression analysis at the TAD level, combining all the genes in a TAD together. It also revealed TAD3189 to have the significantly higher expression from haplotype A among all the TADs on chromosome 8 (Figure 6C). These results are in line with the TAD specific chromatin interactions (from Hi-C and 4C-seq) reported above (Figure 5B, Supp Fig 6A).

A recent study has reported that HPV integrations are enriched in super-enhancers (SE) which ultimately control lineage determining genes (Warburton et al. 2021b). Hence, we asked whether the HPV18 integration in HeLa affects the SE activity on chromosome 8. TAD3189 also harbours multiple SEs, one of which overlaps HPV integration. Interestingly, this SE is the most strongest in terms of signal (enhancer marks) on chromosome 8, and the third most strongest across the genome in HeLa (SEdb (Jiang et al. 2019)). Allele-specific expression analysis using GRO-seq data revealed this SE in TAD3189 to be the most significantly and highly expressed from Haplotype A among all the SEs on chromosome 8 (Figure 6D). Taken together, these results indicate that the HPV integration in chromosome 8 lead to changes in regulatory activity and chromatin interactions, resulting in allele-specific expression of SE and oncogenes within the same TAD (TAD3189).

## Discussion

HPV DNA integration into the host nuclear DNA is often found in the tumours of advanced cervical cancers, as compared to their early stages where the HPV is mostly present in the episomal form (Warburton et al. 2021a; Moody and Laimins 2010; Daniel et al. 1997). Thus, the HPV DNA integration is considered as an important oncogenic event in the transformation and progression of cervical cancer, however, the molecular mechanism underlying this is not yet fully understood. On one hand, it can be attributed to the overexpression of viral oncogenes (E6 and E7) from the integrated HPV DNA which could affect the cellular functions (such as cell proliferation and immune response) (Yim and Park 2005; Rodrigues et al. 2019). On the other hand, the integrated HPV DNA itself can affect expression of nearby host genes in *cis* and thereby contribute to tumour development (Ojesina et al. 2014; Groves and Coleman 2018). However, the extent of the latter at the genome-wide level, particularly in the context of host chromatin structure, is not well understood. Thus, in this study, we attempted to investigate the impact of HPV integration on host chromatin structure and gene regulation genome-wide, by using the HPV integration from cervical tumour samples combined with the chromatin structure information from closest matching cancer and normal cell lines.

First, we showed that the distribution of HPV integrations across the genome is non-random. They are significantly enriched in the active chromatin regions and depleted in the inactive chromatin regions. Though these results are consistent with the previous meta-analysis studies (Doolittle-Hall et al. 2015; Bodelon et al. 2016; Schmitz et al. 2012), our analysis tried to remove any bias emerging from the integration detection methods (such as RNA-seq and APOT) by considering only whole-genome DNA based assays. Additionally, for the enrichment analysis, random integration sites were generated by matching the GC content around the observed HPV integration sites to account for the influence of local sequence content (since the HPV integration processes are associated with microhomology-mediated end joining (MMEJ) (Hu et al. 2015)). This approach is robust as compared to the previous studies which used uniform random distributions. Still the significant enrichment of HPV integration observed at the active chromatin regions in the cervical tumours could be explained by multiple factors acting prior to and/or during the clonal selection. Prior events may include the closer proximity of the episomal HPV DNA to the active chromatin regions due to the interaction between viral E2 protein and host BRD4 chromatin proteins (likely for the utilisation of host transcription/replication factors for viral transcription and replication), DNA breaks at the transcriptionally active regions due to replication-transcription conflicts or torsional stress, DNA repair and microhomology between the HPV and host DNA (Warburton et al. 2021a; Hu et al. 2015; Canela et al. 2017). All these factors can favour the accidental integration of HPV DNA into the host genome. After that, if the host locus of the HPV integration is favourable for the expression of viral oncogenes (E6/E7) and also for upregulation of nearby host genes (especially oncogenes discussed below) that provides the selective growth advantage and favours the clonal expansion, then these integrations are likely to undergo positive selection.

Second, we observed that the transcriptional upregulation in the flanking region (upto 10 kb) of HPV integration was evident in both active and inactive chromatin regions. This could be due to the local genomic alterations (amplifications/translocations) associated with the HPV integration, and also changes in the local chromatin environment. For example, the host transcription factors can bind to the integrated HPV DNA and thereby increase the local chromatin accessibility (Johannsen and Lambert 2013; Karimzadeh et al.; Gagliardi et al. 2020) and subsequently upregulate the expression of nearby host genes. Alternatively, this can be due to the fusion of nearby host genes with the viral genes (Nguyen et al. 2018). Also, it is possible that the HPV integration at host regulatory regions (such as enhancers) could lead to the formation of super-enhancers (Warburton et al. 2018) and thereby enhance the expression of host genes. Our findings expand this observation in cervical cancers as we observed a higher expression of SE eRNAs in the immediate vicinity of the HPV integration regions, regardless of the host chromatin state.

Third, we explored the long-range effect of HPV integration on expression of the host genes. For this, instead of defining arbitrary length cut-offs around the integration sites (Nguyen et al. 2018; Warburton et al. 2021b), we used TAD boundaries as demarcation points. Overall, we found that the TADs with HPV integration showed higher gene expression as compared to samples without integration in the same TADs. And this upregulation is positively correlated with the number of HPV integrations within the TAD. However, genes in the neighbouring TADs were mostly unaffected. This may be due to the fact that the host chromatin structure is influencing the HPV associated genomic alterations (amplification/translocation) during the integration processes or limiting the HPV integration mediated chromatin interaction changes to intra-TAD because of the insulation property of TAD boundaries. Recent studies have shown that HPV integration can cause changes in the local TAD structures in advanced cervical cancers (Cao et al. 2020) and also in human cell lines: HPV16 integration in W12 (prior to clonal selection) (Groves et al. 2021) and HPV16 integration in SiHA cell line (Xu et al. 2021). Our results further extend this observation genome-wide in cervical tumours and show that the majority of the HPV integration induced chromatin changes and associated gene expression changes are mostly confined to the same TAD.

We found few loci with recurrent HPV integration at the TAD level and were associated with overexpression of oncogenes within them. Further, looking at the distribution of HPV integration within these TADs revealed that the expression of oncogenes (such as *MYC* and *PVT1*) were not only affected by HPV integration directly at or closer to these genes, but also by integration farther away (∼500 kb) but within the same TAD. To further understand the mode of regulation of these oncogenes by HPV integration, we chose HeLa as the model system as it had the integration in the most recurrently integrated TAD3189 in cervical tumours. Previous studies have shown chromatin interaction between the integrated HPV DNA and the *MYC* gene (∼500 kb away from the integration) by using ChIA-PET and 3C assays in HeLa (Adey et al. 2013; Shen et al. 2017). However, whether this interaction is localised and specific to MYC regions or the integrated HPV DNA can interact with other genomic regions is not known. Our unbiased analysis, by using available Hi-C data from HeLa, revealed that the majority of the chromatin interactions were localised within the same chromosome, specifically within the same TAD as integration (Figure 5A,B). Further, 4C-seq analysis by taking integrated HPV DNA as a viewpoint, showed haplotype-specific chromatin interaction between integrated HPV DNA and host genomic regions in HeLa. This is further supported by the allele-specific RNA pol II binding enrichment, super enhancer activity and overexpression of *MYC* and *PVT1* genes within the TAD3189. This allele-specific activity can be extrapolated to the cervical tumour samples as well, because TAD3189 is the most recurrently integrated TAD among cervical tumours (Figure 4A), and we also observed the overexpression of oncogenes *MYC* and *PVT1* within them (Figure 4B, C). We propose that this may be one of the many ways by which HPV integration influences the process of tumorigenesis, as role of both of these oncogenes along with the CCAT1 in cervical carcinogenesis have already been established (Shen et al. 2019; Iden et al. 2016; Narisawa-Saito et al. 2012).

The limitations of this study include: (a) The HPV integrations we analysed were mostly detected from the advanced stage cervical tumours, thus their genome-wide distribution with respect to the chromatin states observed here could be influenced by the factors acting prior and during the clonal selection. Future studies which test for *de novo* HPV integration (for example, by using cell line either infected with HPV or genome-wide profiling of early stage cervical tumours) might shed light on the interplay between viral and host factors which drives the integration processes and associated chromatin changes, (b) the overexpression of host regions observed near the HPV integration in active and inactive regions could have contributions from the extrachromosomal DNA (ecDNA), which have a hybrid of viral-host genome (Nguyen et al. 2018). Perhaps in future, the application of long-read DNA/RNA sequencing could help to disentangle the DNA structural conformations of HPV integrations and better quantify the contributions from the ecDNA, (c) the TADs from cell lines HeLa and NHEK covers only 47.5% and 56% of the genome respectively, and so we were limited to analyse those HPV integration that fall within that TAD region. Chromatin interactions maps from the matched tumour samples could help to better understand the long-range effect of HPV integrations genome-wide and also to study the changes of the TAD structure (whether the integration lead to split of existing TAD or merging of adjacent TADs).

To conclude, this study reveals the cis-regulatory potential of HPV integration on the host gene dysregulation through changes in the host chromatin structure and gene-regulatory interactions (Figure 6E). On the basis of our results and previous findings we propose that HPV integration is a strong driver which mediates multiple modes of dysregulation (including overexpression of E6/E7 viral oncogene, cis-regulatory effect of HPV integrations, local copy number changes, and amplification of regulatory elements) that can affect the host cellular functioning and thereby providing selective growth advantages to the cancer cells. This study also demonstrates that TADs can be used to identify host genes at a distance that are likely to be affected by the HPV integrations through looping or chromatin contacts, instead of arbitrary distance cut-offs. This will help to identify more recurrent integrations at the TAD level and also to associate orphan HPV integrations with new target genes. Moreover, our findings reveal the significance of insulated neighbourhoods in the form of TADs and their key role in safeguarding the genome from spurious transcriptional changes driven by viral integrations.

## Materials and Methods

### HPV integrations

We collated HPV integrations in cervical cancer patient samples from previous studies (including TCGA-CESC and others) (Warburton et al. 2021b; Nguyen et al. 2018). This dataset consists of HPV integration identified through genome-wide approaches (whole-genome sequencing [WGS] and HPV capture methods) and exome-wide approaches (whole-exome sequencing [WXS] and RNA-seq based integrations). In total, we got 1324 integrations from 326 samples (see Supplementary Table 1), after removing five samples which had an extreme number of HPV integrations (above the 99th percentile in the respective methods). We categorised the HPV integrations into two main sets: (a) *GW-HPV-int*: containing 617 integrations from 212 samples identified through genome-wide approaches, and (b) *all-HPV-int*: containing 1324 integrations from 326 samples, which includes HPV integrations detected from genome-wide, RNA-seq and exome-based approaches (see Supplementary Table 1). In the latter set, if in any samples, HPV integrations were identified from both WGS and RNA-seq, we merged the overlapping integration to avoid redundancy. For the analysis shown in Figure 1 and 2, we used *GW-HPV-int* set whereas for the others (Figure 3, Supp Figure 3, Figure 4) we used *all-HPV-int* set. Only samples from TCGA-CESC were used whenever the expression was being plotted together with the HPV integration.

### Genomic annotations

ChromHMM regions (15 states) for both HeLa and NHEK cell lines were obtained from the Roadmap Consortium (Roadmap Epigenomics Consortium et al. 2015). We broadly classified these states into two groups: active state (TssA, TssAFlnk, TxFlnk, Tx, TxWk, Enh, EnhG, ZNF/Rpts) and inactive state (Het, TssBiv, BivFlnk, EnhBiv, ReprPC, ReprPCWk, Quies) according to the (Roadmap Epigenomics Consortium et al. 2015).

The individual histone marks dataset used in Figure 1E was obtained from ENCODE (goldenPath/hg19/encodeDCC/).

The genomic coordinates of non-B forms DNA conformations were obtained from http://nonb.abcc.ncifcrf.gov/.

Super-enhancer coordinates for HeLa were obtained from SEdb (http://www.licpathway.net/sedb) (Jiang et al. 2019). W12 super-enhancers were obtained from (Warburton et al. 2021b).

HeLa and NHEK TAD coordinates were obtained from (Rao et al. 2014). Further, the overlapping TADs were merged using bedtools mergeBed (Quinlan and Hall 2010) to obtain non-overlapping TADs (McCole et al. 2018) (Supplementary Table 2).

### Enrichment analysis

For genome-wide enrichment analysis of HPV integration shown in Figure 1, the genomic coordinates of HPV integration sites were extended to a total length of 1 kb size from the centre (such that all HPV integrations have uniform size distribution and to compute the GC content around HPV integration sites). To compute the expected integrations, we randomly sampled an equal number of regions which were of the same length and similar GC content to the observed HPV integrations. For each feature, we calculated the number of observed HPV integrations overlapping with that and compared it against the expected HPV integration counts using the Chi-squared test. The p-values were subjected to the multiple-hypothesis testing correction using Benjamini-Hochberg method (FDR). For ChromHMM annotations, we considered the centre of HPV integrations (and random regions) for the overlap.

### Expression analysis

To check the expression of genes and non-coding elements in the immediate vicinity (Figure 2A, C; Supp Figure 2A, C), we downloaded the pre-computed total RNA expression values in the 10 kb region around the HPV integration regions from (Nguyen et al. 2018). These integration regions were defined by merging HPV integrations that were within 10 kb at the sample level (if any, to avoid overlapping biases), and then a 10 kb flanking region was added on both sides to compute the total RNA expression (Nguyen et al. 2018). In the case of enhancer RNA (eRNA) expression, we followed the above steps, to compute the eRNA expression (from Super-enhancers) around the HPV integration regions. In case of TAD level expression analysis (unique TAD and sample combination), we computed the mean expression using all genes (with expression from TCGA-CESC) in the respective TADs. The eRNA expression from super-enhancer regions and gene expression of TCGA-CESC samples were obtained from the TCeA database (https://bioinformatics.mdanderson.org/public-software/tcea/) and gdc portal (https://portal.gdc.cancer.gov/), respectively.

For the gene level expression comparison with respect to HPV integration status (Figure 4), we used normalised RSEM values. The expression level represented as z-score in Supplementary Figure 4 was calculated by using the mean and standard deviation from all the samples with and without integrations (at gene level).

### Allele-specific analysis

For allele-specific expression/tf-binding analysis GATK ASEReadCounter (v4.1.9.0) was used (Van der Auwera and O’Connor 2020). Only those reads with minimum base quality of 10 and minimum read mapping quality of 20 were used. Also minimum read depth of 8 reads (5 for GRO-seq and TF ChIP-seq datasets) at each heterozygous SNPs was used as a cutoff. Only those features which were supported by at least 15 reads at the heterozygous SNP positions in total were further used. For copy number correction, we used read depth from WGS. Binomial test followed by Bonferroni correction was used to detect the significance of allele specific expression/tf-binding with respect to copy number at each feature level from haplotype A.

For the haplotype-specific binding analysis of RNA polymerase 2 (Pol2) in HeLa, the bam files were obtained from ENCODE (encodeDCC/wgEncodeSydhTfbs/). HeLa RNA-seq data and haplotype specific heterozygous SNP positions from HeLa genome (Adey et al. 2013) were obtained from the dbGaP repository phs000640.v1.p1. The GRO-seq data for HeLa was obtained from GSE63872.

### 4C-seq experiment and data analysis

The 4C-seq experiment was done following the protocol mentioned in (Walavalkar et al. 2020). The primary digestion was performed with DpnII (NEB) and the secondary digestion was done with NlaIII enzyme (NEB). The primer used for the HPV18 viewpoint in HeLa and the number of reads in each replicate is mentioned in the Supplementary Table 3. 4C-ker package (Raviram et al. 2016) was used to analyse the 4C-seq data and the hg19 genome was used as reference.

### Hi-C analysis

Hi-C data from HeLa was obtained from ENCODE (https://www.encodeproject.org/experiments/ENCSR693GXU/). A hybrid hg19-HPV18 genome, considering the HPV18 genome as an additional ‘chromosome’, was constructed (human genome version: GRCh37.75, HPV18 version: NC_001357.1; the HPV18 genome orientation was reversed, as shown in Adey et al. 2013). The hybrid hg19-HPV18 genome was indexed with bwa v0.7.17 ‘index’ mode and samtools v1.6 ‘faidx’. Hi-C reads from each of the replicates were mapped separately to the hybrid genome using bwa (v0.7.17, ‘mem’ mode, parameters: -t 20 -E 50 -L 0 -v 0). Filtering of reads was done based on mapping parameters (min mapq=1, samtools view -Sb -q 1 -F 256). After intersecting reads with HINDIII intervals and removing self ligation and duplicate pairs, replicates were pooled together. HiCExplorer (v2.1.1) was used to obtain the contact list and the contact matrices at 10 kb resolution.

## Acknowledgements

We thank the members of RS and DN lab for feedback and suggestions. The results shown here are in part based upon data generated by the TCGA Research Network: https://www.cancer.gov/tcga. The genome sequence described/used in this research was derived from a HeLa cell line. Henrietta Lacks, and the HeLa cell line that was established from her tumor cells without her knowledge or consent in 1951, have made significant contributions to scientific progress and advances in human health. We are grateful to Henrietta Lacks, now deceased, and to her surviving family members for their contributions to biomedical research. This study was reviewed by the NIH HeLa Genome Data Access Working Group. The genomic datasets used for analysis described in this manuscript were obtained from the database of Genotypes and Phenotypes (dbGaP) through dbGaP accession number phs000640.v1.p1.

## Author contributions

AKS and RS conceived and designed the study. AKS performed most of the data analysis, interpreted the results, and prepared figures. DT and GC contributed for the Hi-C data analysis. KW and DN performed 4C-seq experiment. AKS and RS wrote the manuscript with input from other authors. All authors read and approved the final manuscript.

## Supplementary Figures

**Supplementary Figure 1:**
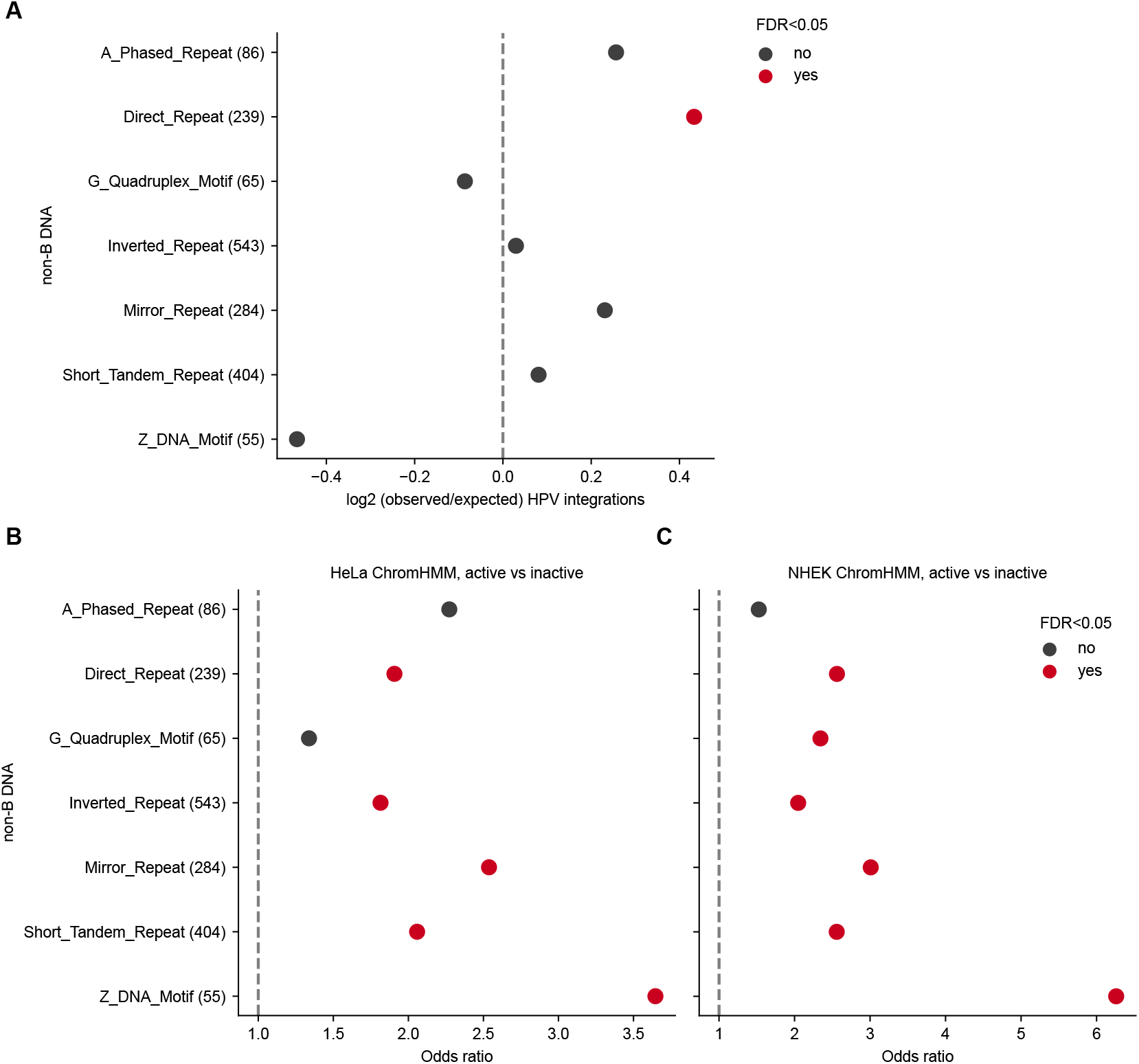
Enrichment of HPV integrations in non-B form DNA regions. **A)** Enrichment of HPV integrations in different non-B forms of DNA predicted genome-wide. The x-axis represents the log2 of observed/expected number of HPV integrations. The y-axis represents the different non-B DNA forms and the observed number of HPV integrations overlapping them (given in the bracket). The p-value was computed using Chi-squared test followed by FDR correction. The colour of the dots indicates whether the adjusted p-value is below the significance level of 5% or not. **B-C)** The enrichment of HPV integrations overlapping each of the non-B forms DNA in active versus inactive regions from HeLa **(B)** and NHEK **(C)** cell lines. The x-axis represents the odds ratio calculated as the ratio of observed versus expected HPV integrations in the active region divided by the ratio of observed versus expected HPV integrations in the inactive region. The p-value was calculated using Fisher’s exact test followed by FDR correction. The colour of the dots indicates whether the adjusted p-value is below the significance level of 5% or not.

**Supplementary Figure 2:**
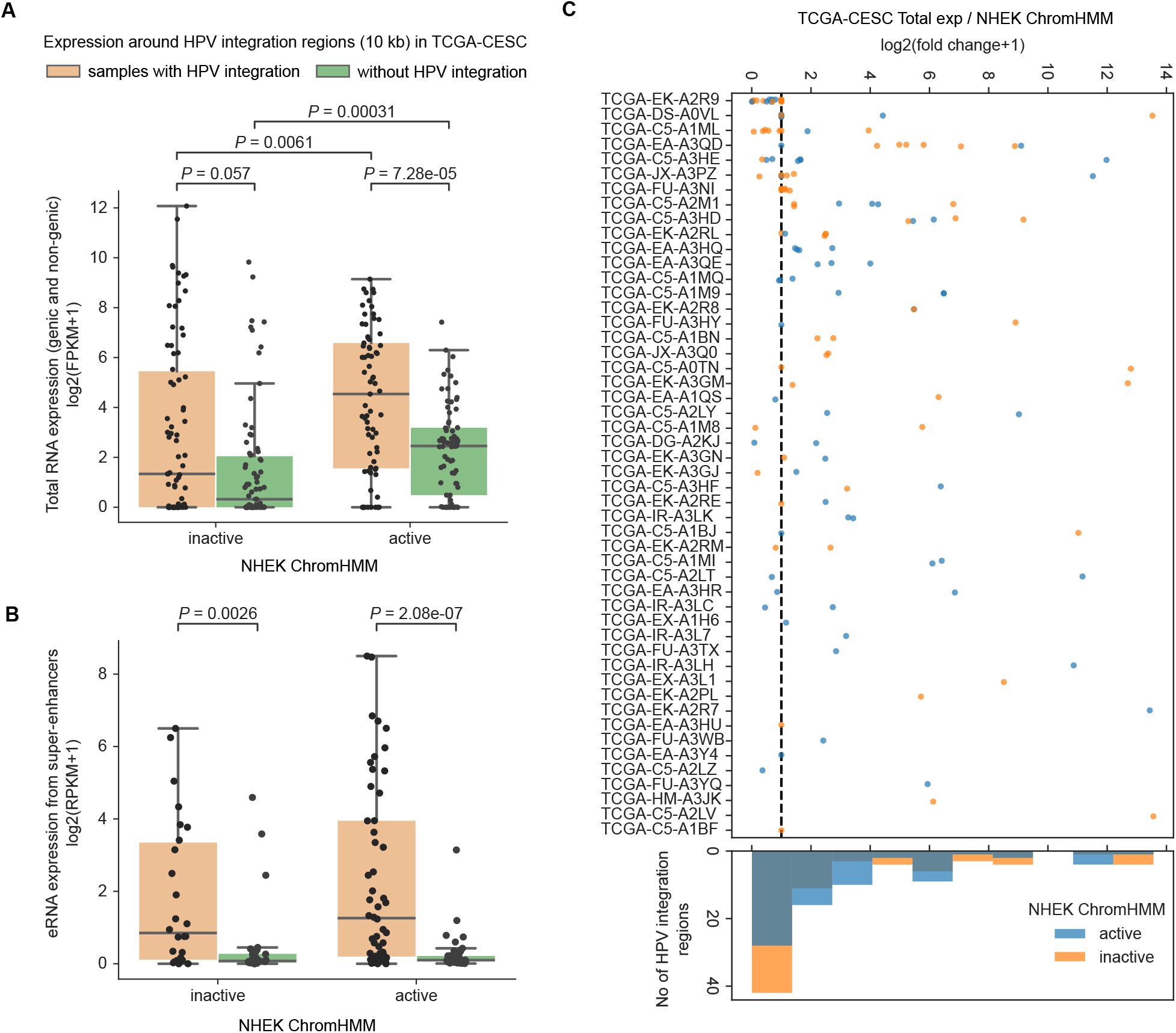
Enhanced transcriptional activity near HPV integration in the context of chromatin states from NHEK. **A)** Boxplot showing the total expression in the 10 kb flanking region around the HPV integration regions as compared to mean expression from TCGA-CESC samples without HPV integration in the same region. The x-axis represents whether the HPV integration is located in an active or inactive chromatin region with respect to NHEK ChromHMM. The p-values were computed using Mann-Whitney U test (two-sided). **B)** Same as **(A)** but for the eRNA expression from super enhancers within 10 kb on either side of the HPV integration regions. **C)** Expression fold change associated with each of the HPV integration regions. The x-axis represents the log2 fold change, which was calculated as the total expression in the 10 kb flanking region around HPV integration regions divided by the mean expression from other samples without HPV integration in the same genomic region. The y-axis represents the individual sample-id of TCGA-CESC samples. The colour of the dots indicates if the integration overlaps an active or inactive ChromHMM region of NHEK. The black vertical line represents the value of log2(fc+1)=1. The histogram at the bottom shows the frequency of integration regions at different fold-change bins. Each dot in **(A-C)** represents a HPV integration region from a sample.

**Supplementary Figure 3:**
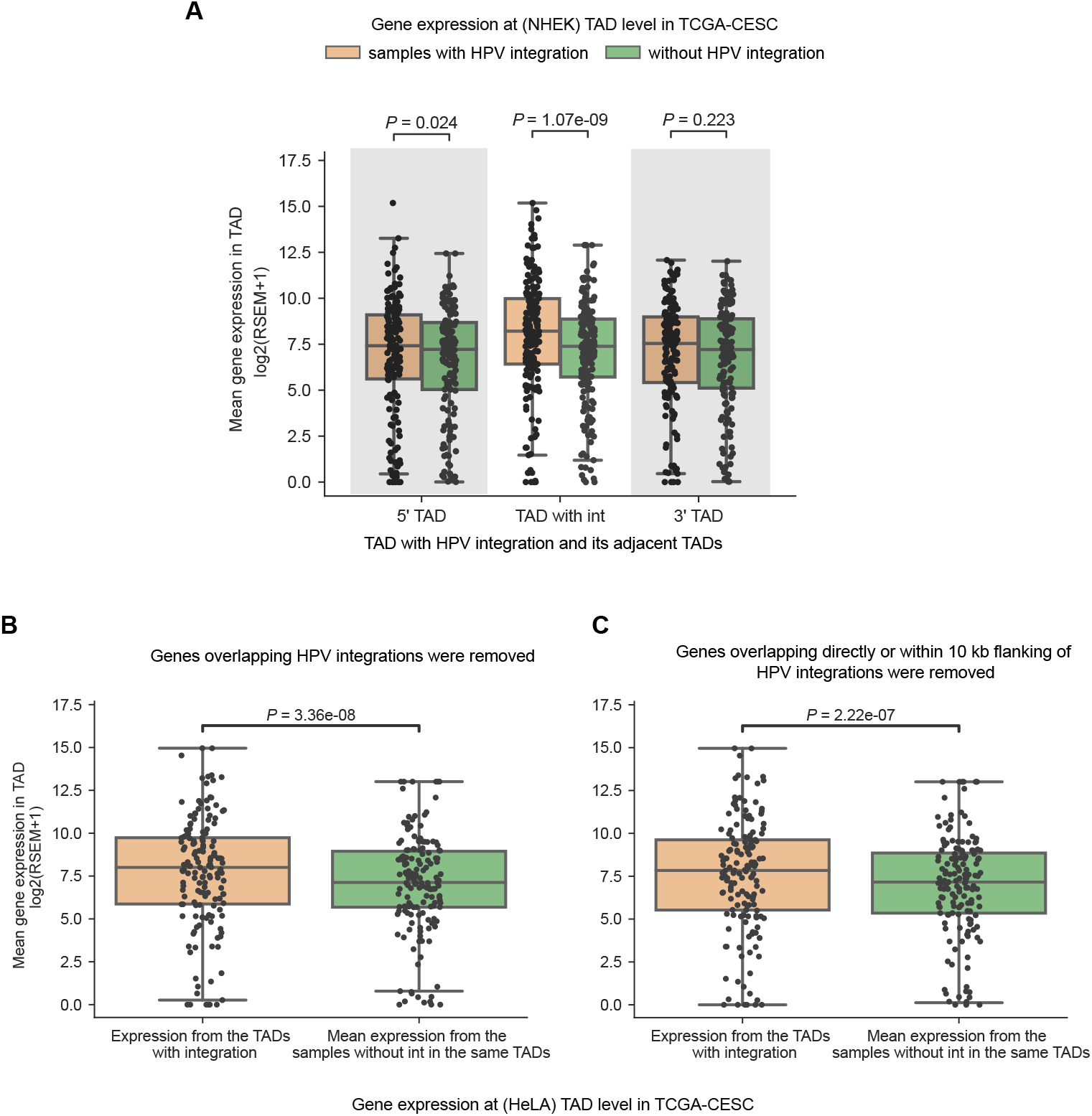
HPV integration associated host gene overexpression with respect to HeLa and NHEK TAD domains. **A)** TAD level gene expression in the TCGA-CESC samples with HPV integration compared to the mean expression from the samples without HPV integration in the same TADs, also for the neighbouring upstream (5’) and downstream (3’) TADs. The TAD information was obtained from the NHEK cell line. **B-C)** TAD level gene expression in the TCGA-CESC samples with HPV integration compared to the mean expression from the samples without HPV integration in the same TADs after removing any genes which were directly overlapping with the HPV integration **(B)** and also after removing any genes which were either directly overlapping or within 10 kb region of an integration **(C)**. The TAD information was obtained from the HeLa cell line. The p-values shown in **(A-C)** were computed using the Wilcoxon signed-rank test (two-sided).

**Supplementary Figure 4:**
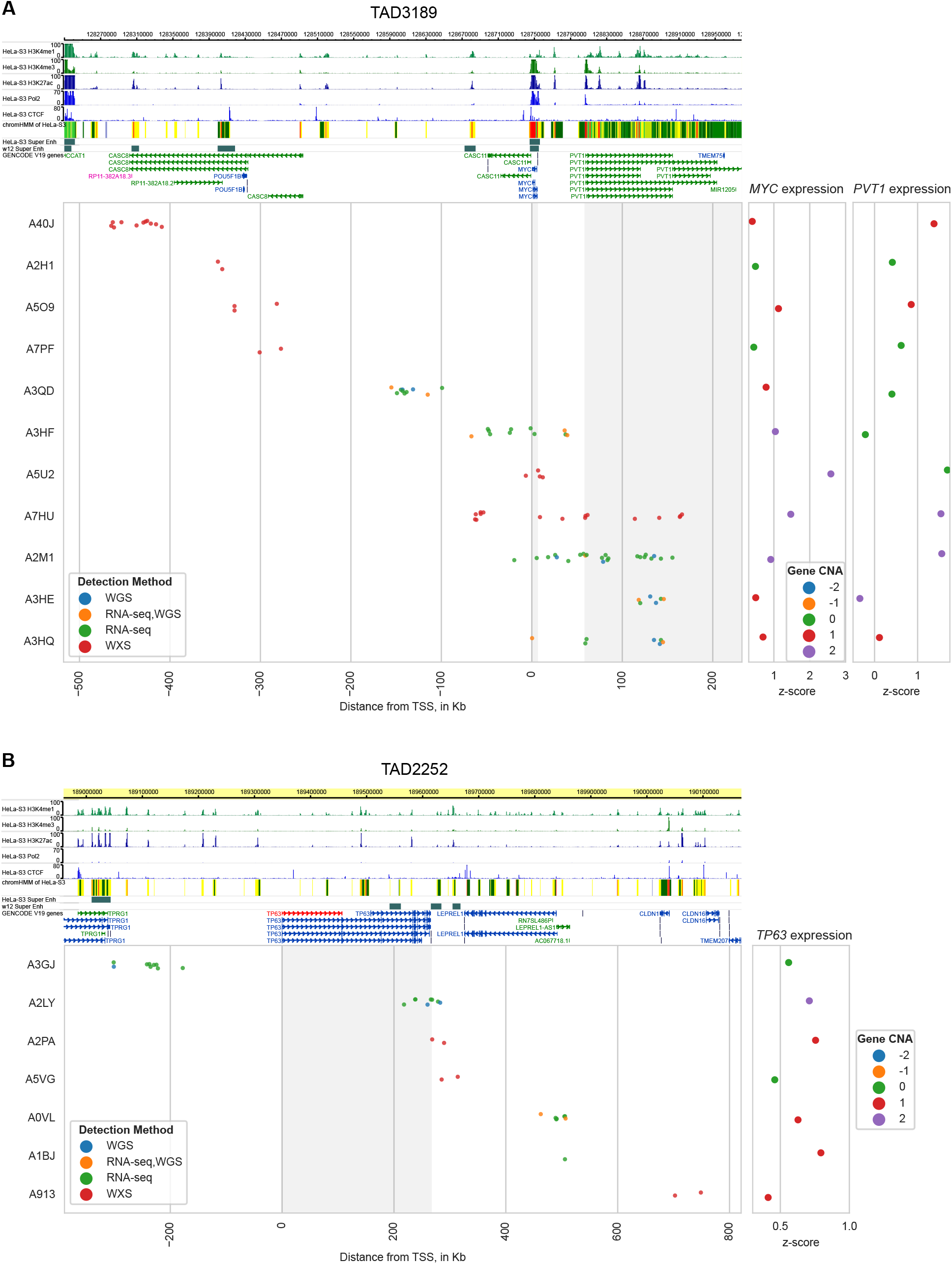
Relationship between distance of HPV integration and the expression of oncogenes. **A-B)** Figure shows all the TCGA-CESC samples with integration in **(A)** TAD3189 and in **(B)** TAD2252. Each of the rows indicates a separate sample with integration (detection method colour coded) in the respective TADs. The MYC and PVT1 gene coordinates falling in TAD3189, and the TP63 gene coordinates falling in TAD2252, are highlighted (in grey). The values on the x axis represent the distance from the promoter of MYC in **(A)** and promoter of TP63 in **(B)**. For each of the samples, the gene expression was represented as z-score on the right-side, and the dots were coloured with respect to relative copy number status of the gene (−2 deep deletion, -1 deletion, 0 copy neutral, 1 amplification, 2 high amplification).

**Supplementary Figure 5:**
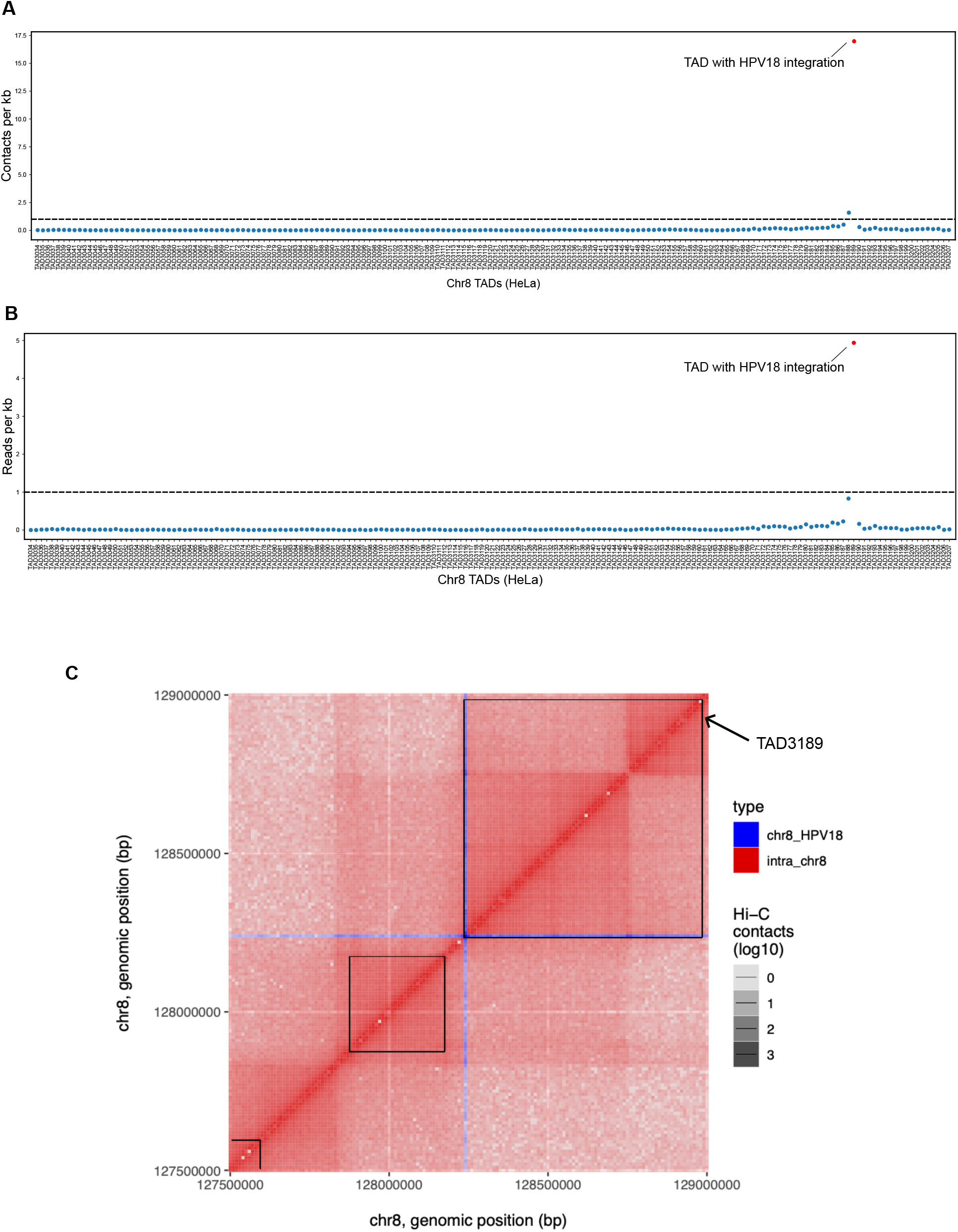
TAD level interaction frequency between HPV18 integrated DNA and host genome in HeLa. **A)** Scatterplot shows the contacts per kb between all the TADs on chromosome 8 and the HPV18 genome. **B)** Scatterplot shows the reads per kb coverage for all the TADs on chromosome 8. These reads represent interaction between that particular TAD and the integrated HPV18 genome. All the TADs are arranged in a linear manner and TAD3189 with HPV18 integration is marked in red in **(A)** and **(B)**. **C)** The heatmap shows the normalised Hi-C contact frequency between integrated HPV18 DNA and chromosome 8 (blue) and intra-chromosome 8 (red) regions. The black square on the top of the heatmap highlights the TADs. The TAD3189 encompasses the majority of interaction between integrated HPV18 DNA and chromosome 8.

**Supplementary Figure 6:**
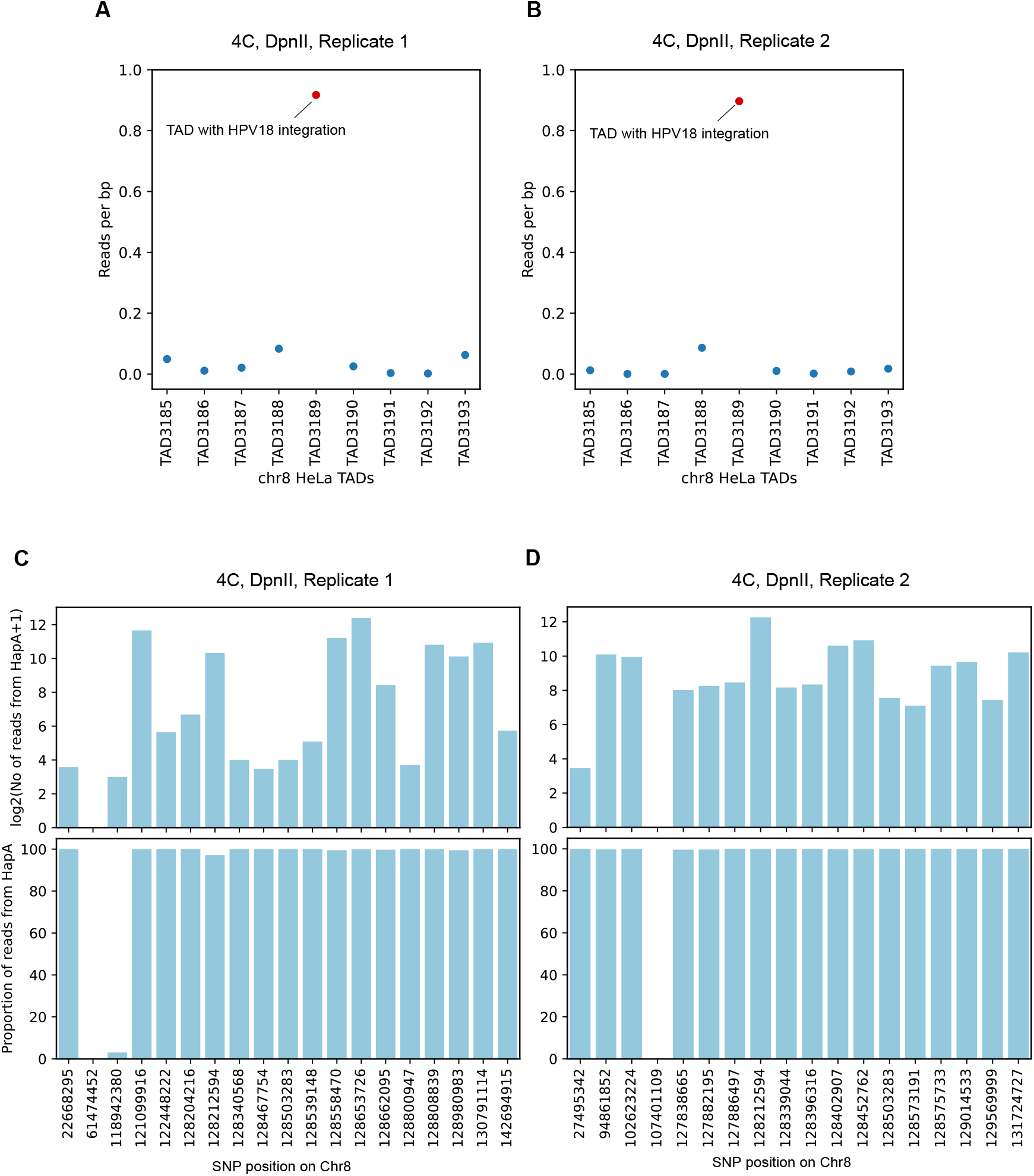
4C-seq analysis reveals haplotype-specific chromatin interactions mediated by integrated HPV18 DNA in HeLa. **A-B)** Scatter plot shows the reads per bp coverage at TAD level, from the 4C-seq experiment for each of the replicates, for TAD3189 and 4 upstream and 4 downstream TADs. **C-D)** Upper bar plot shows the 4C-seq read coverage for the Haplotype A at the heterozygous SNPs which overlaps the 4C-seq coverage regions and lower bar plot shows the proportion of 4C-seq reads mapping to Haplotype A at each of these heterozygous SNPs, for each of the replicates.

## Supplementary Tables

Supplementary Table 1: List of HPV integrations from cervical tumour samples

Supplementary Table 2: HeLa and NHEK TAD co-ordinates.

Supplementary Table 3: Primers used in 4C-seq experiment.

